# Harry Potter meets Markov: Neural event representation in the reading network during narrative processing

**DOI:** 10.1101/2025.10.13.682063

**Authors:** Huidong Xue, Filipp Dokienko, Francesco Gentile, Bernadette M. Jansma

## Abstract

People segment ongoing information into meaningful mental representations known as events. Recent studies have found a spatio-temporal hierarchy in the event structure during movie watching and story listening. How the human brain segments stories while reading remains unclear. We examined event segmentation while reading Harry Potter using publicly available 3T fMRI data. To identify neural events, we employed a Hidden Markov Model (HMM) within six regions of interest (ROI) in the reading network. The ROIs included the inferior frontal gyrus (IFG), middle frontal gyrus (MFG), angular gyrus (AG), inferior frontal gyrus orbital (IFGorb), anterior temporal lobe (ATL), and posterior temporal lobe (PTL). These regions are also associated with DMN. The results revealed multiple timescales of events, with the IFG and AG representing shorter events, while the IFGorb, ATL, PTL and MFG represent relatively longer events. Boundaries of longer events were partially aligned within short events, suggesting a nested hierarchical event structure. Further comparisons identified associations between neural events and annotations from humans and ChatGPT (automated and less subjective segmentation) in the inferior frontal, temporal and parietal areas. Notably, the AG (a core region of the DMN and memory processing) and the IFG (relevant for language processing) exhibited stronger alignment with both sets of annotations. The findings suggest an important role of AG and IFG in event segmentation in reading. They also confirm the connection between the reading network and the DMN in event segmentation. This connection seems to be linked to the hierarchical processing of context-dependent semantic information during reading.

## 1. Introduction

We perceive continuous information from the environment in daily life. According to the event segmentation theory, this information is automatically segmented as discrete events with an identifiable beginning and ending in a specific sequence (Kurby & Zacks, 2008). Event boundaries are associated with multiple changes in stimuli over time, including variations in character, action, time and location, which activate widespread brain regions (Pettijohn & Radvansky, 2016; Radvansky & Copeland, 2010; Zwaan & Radvansky, 1998). A growing body of neuroimaging studies has shown that the neural events could be detected from brain activity, and the default mode network (DMN), which includes the anterior medial prefrontal cortex, medial temporal cortex, angular gyrus, and posterior medial cortex, is sensitive to event changes (Stawarczyk et al., 2021; Wang et al., 2024). When watching movies or listening to stories, our brain represents events in a temporal hierarchical structure, and boundaries across different areas are aligned partially, suggesting a nested hierarchy of event segmentation (Baldassano et al., 2017; De Soares et al., 2024; Hasson et al., 2015; Wolff et al., 2022). Baldassano et al. (2017) asked participants to watch a Sherlock Holmes episode for 50 minutes while brain activity was recorded by fMRI. They found that short events (in seconds) were present in the early visual cortex, and longer events (in minutes) were present in the angular gyrus (AG) and posterior cingulate cortex (PCC). In addition, event boundaries from the AG and PCC aligned with events in the visual cortex, showing a partially nested event structure. Follow-up studies using audiovisual or auditory narrative found relatively low-frequency event segmentation in the prefrontal cortex and PCC. These boundaries exhibited a higher alignment with the events derived from human annotation, suggesting that longer events might be related to the semantic nature of the story (Baldassano et al., 2017; Geerligs et al., 2022; Wu et al., 2025). Furthermore, recent studies have confirmed that the representation of neural events is linked to the function of these areas. For example, Oetringer et al. (2025) revealed that the event boundaries in parahippocampal place area - involved in the processing of visual scenes and locations - closely corresponded to the moments for place changes observed during movie watching (Oetringer et al., 2025). These findings highlight the region’s specialized function in event segmentation within a specific stimulus context. However, how the event segmentation is organized within the reading network remains elusive.

When reading a story, a reader integrates general knowledge and a rich narrative context to identify event boundaries from a sequence of words. Several behavioral studies in narrative reading suggested that event segmentation temporally aligns with changes in situation features, such as time shifts, change of spatial location, and characters. Changes across events seem to elongate reading duration (Pettijohn & Radvansky, 2016; Swets & Kurby, 2016; Zacks et al., 2009; Zwaan et al., 1998). Situational segmentation seems directly related to how we memorize and recall information, and that time around event boundaries is crucial for it (Bernhard et al., 2024). The neuroimaging results also showed increased brain activity in readers after a situational event boundary (Speer et al., 2009; Speer et al., 2007). Similar to findings for movie watching, different situational features in story reading activate distinct brain regions. For example, the introduction of a new character was associated with increased activity in the bilateral posterior superior temporal cortex and superior frontal gyrus, while changes in the character’s action goals were linked to increased activity in the middle frontal gyrus (Speer et al., 2009). These regions belong to the dorsal pathway of language processing (Fridriksson et al., 2016; Woolnough et al., 2023). Additionally, various kinds of descriptions within the story activated different regions of the brain during reading. Motor descriptions (concrete actions, e.g. *They reached the bus-stop shelter*) elicited activity in the cingulate cortex, superior frontal gyrus, and middle frontal gyrus, which are components of the dorsal pathway (Marek & Dosenbach, 2018). Mental event descriptions (reflecting thoughts of the protagonist, e.g. *She thought of the recurrent waves of pain*) were represented in the temporal pole, parietal operculum, anterior cingulate, and angular gyrus within the ventral pathway (Mak et al., 2023; Marek & Dosenbach, 2018).

In the context of reading, the ventral pathway is devoted to semantic processing, and the dorsal pathway is involved in phonological processing (Friederici & Gierhan, 2013; Goranskaya et al., 2016; Kearns et al., 2019; Saur et al., 2008; Woolnough et al., 2023). Additionally, accumulating evidence suggests that the dorsal pathway supports event construction, while the ventral pathway supports event detection and maintenance, and the core of the DMN underlies event representation and prediction (Sridharan et al., 2007; Stawarczyk et al., 2021; Yarkoni et al., 2008). Taken together, the DMN is primarily proposed to support event segmentation, while both the ventral and dorsal pathways contribute to language processing and event segmentation. The DMN and language regions might share fundamental neural event segmentation mechanisms during narrative processing in movie watching and story listening. In this context, the reading network is hypothesized to perform event segmentation in a similar hierarchical nesting structure. Notably, regions that highly overlap with DMN should exhibit event segmentation patterns similar to semantic event annotations from humans and larger language models.

In this study, we aimed to address these questions about neural events represented in the dorsal and ventral pathways of the reading network. We do so by studying neural event segmentation during natural reading within pre-specified ROIs of the reading network, which spans both the dorsal and ventral pathways of the reading network and the DMN. We examined brain activity with a publicly available fMRI dataset in which participants read chapter nine of “Harry Potter and the Sorcerer’s Stone” (Wehbe et al., 2014). This dataset is a key resource and has been widely used in numerous studies, providing valuable insights into reading processing (Hamilton & Huth, 2020; Toneva et al., 2022; Xiong & Cribben, 2022). To reveal the neural event structures within cortical regions, we adopted a Hidden Markov Model (HMM) based data-driven approach to identify temporal transitions between neural events (Baldassano et al., 2017). The approach entails two assumptions. The first is that the readers perceive a sequence of discrete event representations (neural events/ latent states) while processing a narrative. The second is that each neural event has a distinct signature (a multi-voxel fMRI pattern representation) that corresponds to the event structure in the narrative. Utilizing the HMM, we identified the transition time points from one event to another in the fMRI time course data, based on the probability distribution of transfers. Because the model can be fitted with different event counts, we can estimate each region’s preferred timescale of event representation by their optimal number of events. We use the t-distance (TD) metric to identify this optimal count, which maximizes correlations of time points within the same events while minimizing correlations of time points within consecutive events (Geerligs et al., 2021). This approach enables us to determine optimal event structures in reading-related regions.

To interpret the HMM solution well, most studies compare neural event boundaries with human annotation (Baldassano et al., 2017; Geerligs et al., 2022; Wu et al., 2025). Positive correlations then support the validity of detected neural events. In addition, emerging large language models (e.g., Generative Pre-trained Transformer Networks, GPT) can serve as the source for event segmentation in narratives. Studies have found that event structure generated by GPT resembled a human-like pattern and exhibited low variation between iterations (Kumar et al., 2023; Michelmann et al., 2025; Panela et al., 2025). Unlike humans, GPT can automatically segment events from text with less subjectivity. Therefore, we compared the temporal alignment of HMM-derived neural event boundaries per region to those annotated by humans from a separate group of volunteers, as well as to the annotations provided by GPT. Any alignment between HMM-derived event boundaries within a region and annotations would reveal its relevant role in segmenting content during reading, reflecting the shared temporo-spatial dynamics of neural events in both reading and event segmentation (Northoff et al., 2025).

## 2. Methods

### 2.1. Event segmentation for neuroimaging data

#### 2.1.1. Harry Potter fMRI dataset

The fMRI dataset used in the present study included eight subjects reading chapter nine of “Harry Potter and the Sorcerer’s Stone” (Wehbe et al., 2014). The chapter consists of approximately 5000 words that were presented in four runs by rapid serial visual presentation (RSVP), and each word appeared for 0.5 seconds at the center of the display (Fig. 1A). Participants were asked to read the story silently. Imaging data were acquired using a T2*-weighted echo planar imaging pulse sequence (repetition time (TR): 2s, echo time: 29 ms, flip angle: 79°, 36 slices, voxel size: 3 × 3 × 3 mm). The anatomical data was acquired using a T1-weighted 3D-MPRAGE pulse sequence. The dataset was collected by Wehbe et al. from nine subjects, but data from one subject were excluded due to excessive noise, leaving data from eight subjects available for use. The preprocessed version of this dataset was obtained from a recent study, with smoothing, detrending, and artifact removal applied, and 1291 TRs remaining for analysis (Reddy & Wehbe, 2021).

**Fig. 1.**
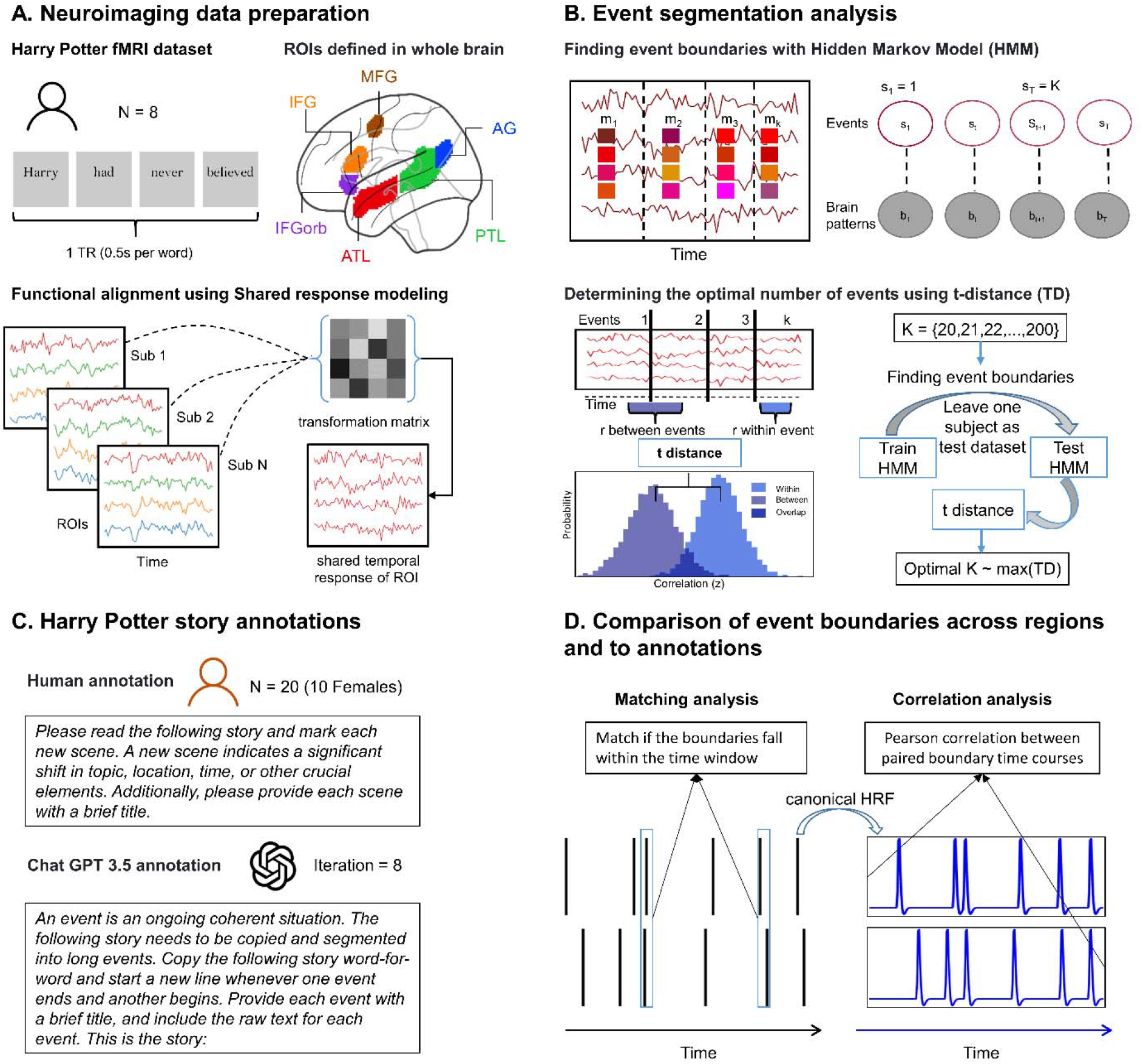
This diagram summarizes the data collection and analysis process in four stages, from neuroimaging data processing to event segmentation, annotation collection, and event boundaries comparison. (A) A publicly available 3T fMRI dataset acquired from 8 subjects reading chapter 9 of “Harry Potter and the Sorcerer’s Stone”. Six ROIs were defined in both hemispheres: inferior frontal gyrus (IFG), middle frontal gyrus (MFG), angular gyrus (AG), inferior frontal gyrus orbital (IFGorb), posterior temporal lobe (PTL), and anterior temporal lobe (ATL). The fMRI data was functionally aligned across subjects for each region using shared response modelling, which estimated the subject-specific transformation matrix and produced a shared temporal response. (B) HMM infers the latent state S_t_ for each time point of the shared temporal response time course per ROI and by doing so, it denotes the event to which that time point belongs, starting in event 1 and ending in event K. All data points during event k are assumed to exhibit high similarity with an event-specific pattern m_k_. Time points belonging to the same event should be more similar, and those within adjacent events should be less similar. The t-distance distinguishes this similarity by calculating the distributions of Pearson correlation between consecutive events and those within the same event. The optimal number of events (k) was determined by the maximum t-distance (TD) within a range of 20 to 200 k. (C) Twenty subjects were asked to segment the Harry Potter story into events with a short title. The same task was performed by Chat GPT 3.5 (GPT-3). (D) Matching analysis: we measured boundary alignment across regions and among neural, human and GPT consensus annotations by calculating the fraction of top-level boundaries near a boundary in the bottom level. “Near” was defined as within a time window of 3TRs (represented by the blue box), with the box sliding across the entire time course. Correlation analysis: we applied the canonical HRF algorithm to the original annotation boundary series (proportion of participants or GPT-3 iterations that marked event boundaries per time point). We then correlated it with the HMM-generated event probability time course by Pearson correlation.

#### 2.1.2. ROIs in the reading network

We focused on six regions within the reading network, spread across the dorsal and ventral pathways, including the inferior frontal gyrus (IFG), middle frontal gyrus (MFG), angular gyrus (AG), inferior frontal gyrus orbital (IFGorb), anterior temporal lobe (ATL), and posterior temporal lobe (PTL). These predefined regions of interest (ROIs) in this dataset were defined using an established language network mask based on a contrast between sentence, word list and non-word sequence reading (i.e. Jabberwocky, non-word list) (Fedorenko et al., 2010). The regions were initially identified in the left hemisphere and then mirrored to the right hemisphere (Fig. 1A). According to the previous studies, we merged the voxels from both hemispheres for each region (Baldassano et al., 2017).

#### 2.1.3. Functional alignment with shared response model

To align fMRI data and capture reliable brain activity evoked by the story across participants, we used the shared response model (SRM) to align all subjects’ data into a shared multi-dimensional space for each ROI. SRM can resolve functional-topographical idiosyncrasies across subjects by identifying a common, low-dimensional feature space, where corresponding time points from the same story are positioned closely together (Chen et al., 2015). Given a voxel by time matrix of fMRI data for each region, SRM produces a set of weight matrices of subject specific topographies (voxels by features matrices), and a shared latent temporal response (features by time matrices) by a probabilistic latent factor model (Fig. 1A). Using these transformation matrices, we obtained the shared response for each subject in the SRM space, which we then used to perform event segmentation analysis.

#### 2.1.4. Hidden Markov based event segmentation model

We next captured the signature of each neural event from the continuous shared response using HMM. This approach assumes that stimuli will generate a sequence of discrete brain states in the observers, referred to as neural events. It also assumes that each event in the story corresponds to a unique signature in the observed brain activity. Therefore, using the shared response of fMRI data as the observed data, we could estimate the latent neural events by identifying the stable brain activity patterns during narrative reading (Fig. 1B). The HMM-based event segmentation model specifies the first and last time point of the signal as belonging to the first and last neural event, respectively. All the other timepoints can either stay within the same event as the previous one or shift to the next event. HMM creates a probability distribution for each time point that differentiates “between” and “within” event types. Therefore, for a given brain observation (*b*_*t*_, t is the time point) and event-specific pattern (*m*_*k*_), the probability of the latent state *s*_*t*_ belonging to event K can be modeled using an isotropic Gaussian model: 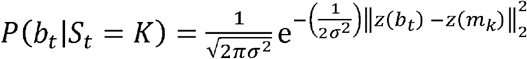. Thus, the log probability of z-scoring brain observations *z*(*b*_*t*_) in the event-specific pattern *m*_*k*_ is a proportion of the Pearson correlation between *b*_*t*_ and *m*_*k*_ plus a constant offset: log *p*(*b*_*t*_|*s*_*t*_ = *k*) ∝ *r* (*b*_*t*_ *m*_*k*_)+ *c*.

To estimate the event-specific pattern *m*_*k*_ and the event structure *s*_*t*_ = *K*, an annealed version of the Baum-Welch algorithm was used. This algorithm is a dynamic programming approach to tuning parameters *m*_*k*_ and *s*_*t*_ by iterative estimation. In the beginning, the setting of distribution *p*(*s*_*t*_ = *K*) is rooted in an assumption that all events have an equal length that leads to the 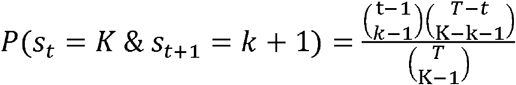 where 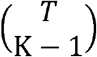 is the placements of event boundaries. Then the event-specific pattern *m*_*k*_ can be calculated as a weighted average 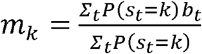, and time points *b*_*t*_ have high similarity with the *m*_*k*_. Subsequently, a new distribution was created based on this *m*_*k*_. However, the accurate event segmentation might include events of different lengths, which is very far from the original assumption of equal event length. To solve this problem, a split and merge algorithm was applied (Baldassano, 2020). This algorithm double-checks whether one event can be better defined as two (split), or re-allocated, merged with adjacent events, in case of higher similarity with the adjacent event (Baldassano, 2020). This function improves the model sensitivity for events differing in length and keeps the same number of events during iteration. For this reason, we applied a split-merge Hidden Markov Model (sm-HMM) via the BrainIAK software (Kumar et al., 2021) to detect neural event boundaries and characterize the event structure in the reading network.

#### 2.1.5. Determining the optimal event count

Although the HMM method can detect latent event boundaries in fMRI data, it is crucial to determine the optimal number of events for each brain area. In this regard, the most common approach is to set a varying number of events, which in this case range from 20 to 200 events, and to find the best event structure (Fig. 1B). According to the event segmentation theory, the best event structure is characterized by a higher correlation of brain response within compared to across events. Therefore, for each solution, the correlations for all paired points within the same and cross consecutive events were calculated, as well as the difference of their distribution using the t-distance (t value, based on t-statistic test) (Geerligs et al., 2021). We used the leave-one-out cross-validation method to train the HMM on seven subjects and test it on the remaining subject for each ROI. We then averaged the t-distance across tested subjects for each number of events. Finally, the solution with the number of events (k) yielding the maximum t-distance was set as the optimal number of events.

### 2.2. Annotations from human and GPT

As the original dataset did not include annotations, we collected human behavioral annotations from 20 volunteers (English level at least B2 of Common European Framework of Reference; 10 females; mean age = 22 years, range: 19-38 years). They were instructed to read the chapter and to highlight changes in character, location, time, or new scenes. To facilitate segmentation into meaningful events, they should also generate a short title for each event (Fig. 1C).

In addition to human annotation of the Harry Potter story, we employed ChatGPT-3.5 (hereafter referred to as GPT-3) for automated event annotation (Chang et al., 2024). To this end, the plain text was input into the GPT-3 event (Fig. 1C). To find an event segmentation, the prompt for the model was set as “An event is an ongoing, coherent situation. The following story needs to be copied and segmented into long events. Copy the following story word-for-word and start a new line whenever one event ends, and another begins. Provide each event with a brief title and include the raw text for each event. This is the story:”. Due to the input limitation of GPT-3, the story was divided into 4 parts with around 200 words of overlap to ensure enough context for event segmentation in GPT-3. The chat was conducted eight times, and the results of the detected boundaries were averaged.

All event boundaries from annotations were marked after complete sentences and aligned to the fMRI data at the TR time scale. To improve comparability to the HMM results, the boundaries close to the previous boundaries within 4 TRs were merged, resulting in the original annotations. Since the participants for annotation were required to generate meaningful events, a boundary supported by four or more individuals was considered a consensus annotation for humans. Consensus boundaries for GPT-3 were based on occurrences of more than two. These original and consensus annotations were used for further comparison analysis.

### 2.3. Comparison of event boundaries across ROIs and to annotations

#### 2.3.1 Matching analysis of event boundaries across ROIs

With the optimal number of events for each region, we trained the sm-HMM for the group dataset and identified neural event boundaries. To quantify the structure of these boundaries across regions, we used a matching analysis. This approach measures agreement between HMM-generated boundary arrays by considering boundaries from two regions as congruent if they fall within a predetermined sliding temporal window (e.g., 3TR) (Fig. 1D). The match fraction was calculated by dividing the number of matched boundaries by the total number of event boundaries. Subsequently, permutation tests were conducted to determine whether the observed match fraction significantly exceeded chance levels. Corresponding null distributions were generated from 1,000 iterations of event order scrambling for one region. This methodological framework enabled exploring potential hierarchical structures within the neural network associated with narrative processing.

#### 2.3.2 Matching analysis for alignment

To evaluate the concordance between boundaries derived from the HMM and human annotations, we used the same matching analysis to estimate the similarity between HMM-derived event boundaries and consensus annotations. Given that the optimal number of events identified for each ROI differed from the annotated counts, potentially affecting the comparison of matching fractions across regions, we refitted the HMM model for each region by using the number of events from human and GPT-3 annotation, respectively. This adjustment helped to reduce confounding effects arising from temporal inconsistencies. Since the annotation has already been adjusted for deviation on the TR scale, we changed the predetermined temporal window to 1 TR. Afterwards, permutation tests were deployed to ascertain the significance of match fractions. We further compared the differences in match fractions across regions to examine how matching with annotation varies by region. For each region, we calculated the matching fraction between scramble HMM boundaries and annotation to create a null distribution against which we assessed the significance of the observed difference.

#### 2.3.3 Correlation analysis for alignment

To validate our results from matching analysis and extending our findings with the optimal number of events, we also assessed the congruence between the optimal event structure in each region and the original annotations from all individuals via a correlation analysis (Fig. 1D) (Yates et al., 2022). To establish robust estimates of the optimal event boundaries occurring across our fMRI sample, we conducted 1000 HMM iterations, resampling participants with replacement, and refitting the model each time. The event probability time course was averaged across all iterations. The outcome of the HMM event time course was then compared to the time courses of human and GPT-3 annotations. To create a continuous boundary time course for the original annotations from humans, for each time point (TR), the amount of observed event boundaries was calculated, and then convolved with a canonical HRF to align with the HMM-derived event boundaries time course based on fMRI data. The same way was also used for GPT-3 annotation. We computed the Pearson correlation between paired time courses of the paired probability of event boundaries. Significance was determined by generating a null distribution through time-shifting the HMM-derived event probability matrix, with wrapping the last *t* time points to the beginning. The value of *t* varied between 1 and the length of the fMRI data. The real correlation values were then compared to the correlations between annotations and the null distributions.

## 3. Results

### 3.1. Event structure in the reading network

To evaluate the optimal neural event structures across the dorsal and ventral pathways for reading, we recognized the maximum t-distance for six regions and compared the event structure with the corresponding null distributions generated by applying HMM to randomly shuffled event order. The analysis revealed that the angular gyrus (AG) and inferior frontal gyrus (IFG) encoded relatively shorter events at a time resolution below one minute, resulting in 67 and 62 events over the entire book chapter reading, respectively. In contrast, the middle frontal gyrus (MFG) encoded relatively longer events, with an optimal event number of 37. Meanwhile, the inferior frontal gyrus orbital (IFGorb), anterior temporal lobe (ATL), and posterior temporal lobe (PTL) encoded events of longer duration, at the minute level, resulting in 44, 45, and 50 events, respectively (Fig. 2A & Fig. 3A). The HMM detected 37 (MFG) to 67 (AG) events across ROI (Fig. 2A), and their boundaries were significantly aligned with each other such that some boundaries in longer events aligned with boundaries in shorter events across regions (Fig. 2B & Fig. 3B). In addition, the adjacent regions (e.g. IFG-IFGorb, IFGorb-ATL, and PTL-AG) showed more matched boundaries. These findings suggest a nested neural event structure within the reading network, with the dorsal and ventral pathways exhibiting hierarchical timescales for neural events (Fig. 2C & Fig. 3B).

**Fig. 2.**
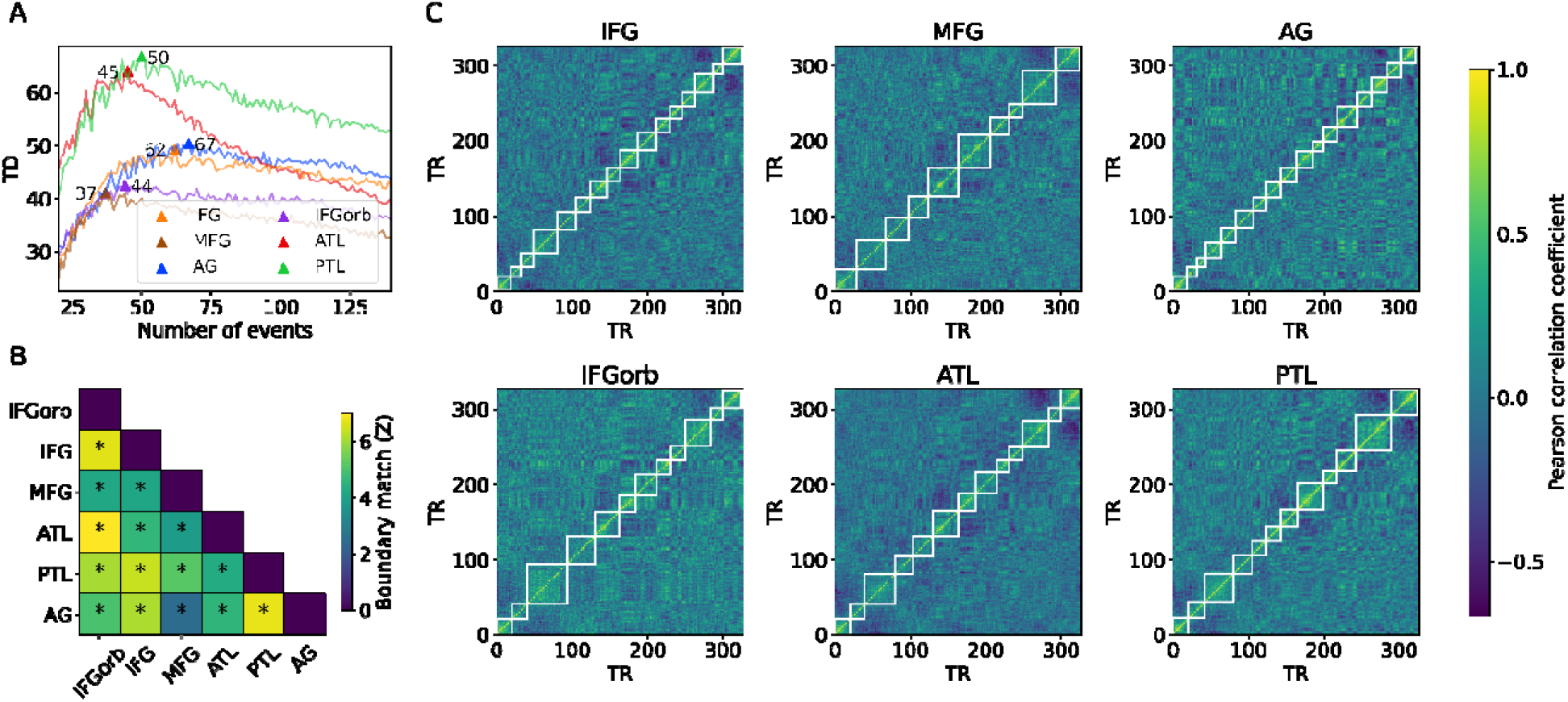
(A) T-distance (TD) reflects the similarity difference of the time points within and across events. A maximum in TD is a metric for identifying the optimal number of events. This resulted in 44-67 events across ROIs (optimal k is the number after the region label): IFG-62, MFG-37, AG-67, IFGorb-44, ATL-45, and PTL-50. (B) The paired matching results for event boundaries showed alignment of boundaries across ROIs within the reading network (yellow = higher alignment). (C) Example TR by TR correlation matrices (for the first 300 TRs) for each ROI in the reading network, with their HMM detected boundaries shown in a white box.

**Fig. 3.**
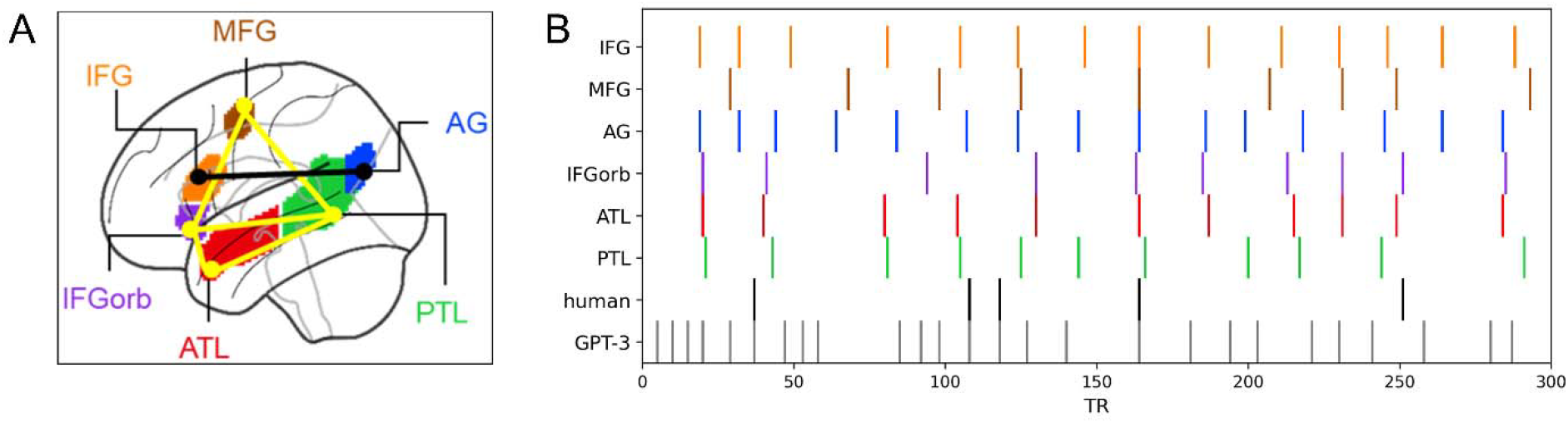
(A) The visualization highlights regions with a similar event structure. Black: IFG and AG, representing relatively shorter events. Yellow: ATL, PTL, IFGorb, and MFG, representing relatively longer event structures during story reading. (B) Visualization of event boundary alignments of HMM, human, and GPT-3 annotation (first 300 TRs as an example). Top 6 lines: ROI boundaries with the optimal number of events for each region over time (TRs), detected by HMM. Line 7-8: consensus annotations from humans and GTP-3 were calculated based on at least 20% agreement from the participants and iterations, respectively.

### 3.2. Events alignment between HMM and annotations

Humans and GPT-3 both identify boundaries for meaningful events from the story of Harry Potter (Fig. 3B). The shared event boundaries were defined when over 20% of human or GPT-3 annotations identified them. Based on this definition, 36 shared events were identified from 81 events in human annotations (M = 17.25; SD = 8.12), and 75 shared events were identified from 109 events in GPT-3 (M = 36.75; SD = 3.60).

#### 3.2.1. Matching analysis results

Using the number of events from human-consented annotation and GPT-3-consented annotation, the HMM was re-fitted for each region, respectively. The match analysis between HMM-derived event structures showed a significant alignment with annotations. Four regions showed significant alignment with human annotations (IFG, *p* = 0.026; AG, *p* = 0.004; IFGorb, *p* = 0.023; ATL, *p* = 0.021; PTL, *p* = 0.213; MFG, *p* = 0.433; Fig. 4A). Further pairwise comparisons show that the AG had larger Human-HMM matches than the MFG (*p* = 0.035) (Fig. 4A). When considering GPT-3 annotation, results revealed significant matches for the IFGorb (*p* < 0.05) and a trend for the IFG (*p* = 0.080) (Fig. 4B). The IFGorb also showed a tendency for larger GPT-3-HMM matches than the ATL (*p* = 0.055) (Fig. 4B). These findings suggest that the IFG, AG, ATL and IFGorb were involved in higher-order segmentation of conceptual events. It shows a combined contribution of the dorsal and ventral pathways to conceptual event segmentation.

**Fig. 4.**
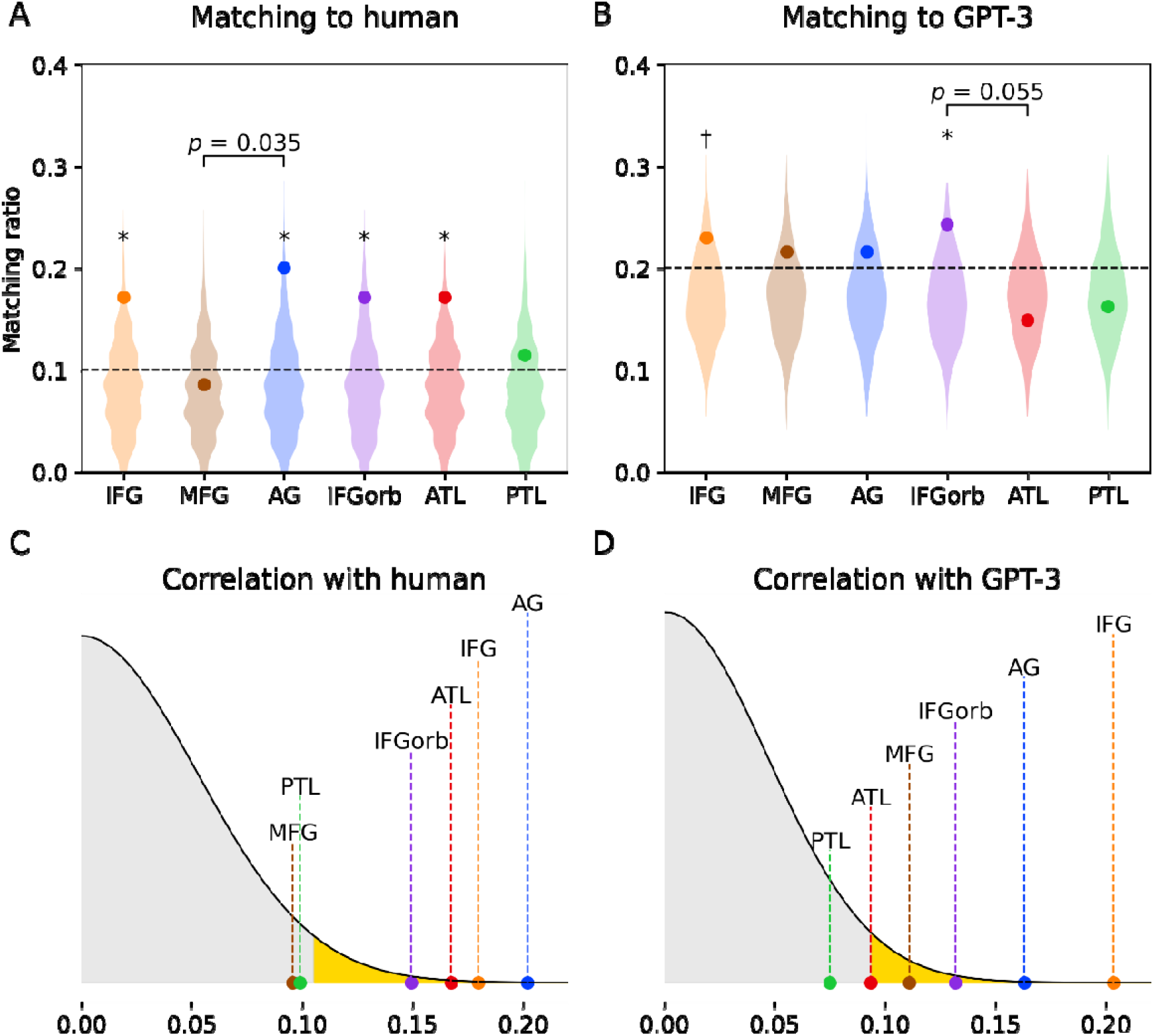
Statistical comparison of HMM-derived boundaries to human and GPT annotation. Upper panel: Matching analysis between HMM-derived boundaries for each region and human annotation (A) and GPT-3 (B) annotation. Each single dot represents the actual matching fraction, while the violin plot depicts the permuted null distribution resulting from time shifts of the neural event boundary time courses. * Indicates p < 0.05; † indicates p = 0.08. Lower panel: Correlation analysis between the optimal event structure in each region and annotations of human (C), and GPT-3 (D). The x-axis represents correlation values (r), and the y-axis represents probability. Single dots represent the actual correlation values, with significant areas highlighted in yellow at the 0.05 level.

#### 3.2.2. Correlation analysis results

Next to matching, we obtained HMM-derived boundary time courses based on the optimal number of events for each region and examined the Pearson correlations between these boundaries and the boundaries identified in the annotations. The analysis revealed significant correlation for neural events and human annotation in the IFG (*r* = 0.185, *p* < 0.001), AG (*r* = 0.201, *p* < 0.001), IFGorb (*r* = 0.152, *p* = 0.020), and ATL (*r* = 0.165, *p* = 0.040). This was not the case for other ROIs (MFG: *r* = 0.094, *p* = 0.140; PTL: *r* = 0.099, *p* = 0.100). The correlation analysis verifies that human boundaries are related to the optimal neural event structure in these regions (Fig. 4C). The analysis also revealed correlation between neural event structure and GPT-3 annotation in IFG (*r* = 0.206, *p* < 0.001), MFG (*r* = 0.110, *p* < 0.05), AG (*r* = 0.162, *p* < 0.001), IFGorb (*r* = 0.129, *p* < 0.001), but not for ATL (*r* = 0.092, *p* = 0. 140) and PTL (*r* = 0.075, *p* = 0.180) (Fig. 4D).

## 4. Discussion

We aimed to establish proof of concept for neural event segmentation in the reading network during story reading. We did so by applying a data-driven method, HMM. To validate the HMM solution of event boundaries, we compared them with humans and GPT-3 annotations. Compared to humans, GPT-3 can be seen as less subjective. We found that the HMM was able to extract event boundaries from the fMRI time course, providing the optimal number of events for each target region. These numbers varied slightly across regions in the reading network, revealing the longest events in the MFG, relatively long events in the PTL, ATL and IFGorb, and the shortest events in the IFG and AG. This HMM-based analysis showed that reading seems to trigger event segmentation in a hierarchical and nested manner, similarly to what has been suggested for movie watching and auditory narrative processing. In addition, the neural events in the frontal, parietal and anterior temporal regions were related to annotations from humans and GPT-3. These correlations validated the HMM results for specific regions within the reading network and confirmed the relevance of these regions for event segmentation during reading.

### 4.1 The hierarchical event structure in the reading network

Our findings suggest the presence of neural representation of events during reading. These neural event representations seem to operate at multiple timescales across regions within the reading network. Notably, events on shorter timescales partially align with those on longer timescales, resulting in a hierarchical structure. This hierarchical event structure is consistent with previous studies on movie watching and story listening. Specifically, short events were observed in sensory regions, while relatively longer events occur in the default mode network (Baldassano et al., 2017; Geerligs et al., 2022; Wu et al., 2025). Our research tested this hierarchical processing within the reading network. This network is crucial for processing language information derived from written words, including the transfer from orthography to phonology, as well as for syntactic and semantic processing of the text. For instance, the IFG, MFG, and AG, within the dorsal pathway, play a role in phonological information processing, both in comprehension and production. The frontal (e.g. IFGorb), temporal (e.g. ATL and PTL), and parietal areas (e.g. AG) are recognized for their involvement in semantic and syntactic comprehension (Chang et al., 2015; Friederici & Gierhan, 2013). Additionally, these regions are involved in the situational dimensions of narrative comprehension (Vaccaro et al., 2021; Xu et al., 2021; Yarkoni et al., 2008; Youssofzadeh et al., 2022). The observed neural event pattern in the reading network reflects a hierarchical process of information integration within these regions, where each region may have a distinct role in event segmentation.

We observed the IFG and AG segmenting events at a relatively high frequency for language processing. We suggest that this reflects the updating and integration of the incoming information into the story context. The results for the IFG align with studies reporting its relevance in fast and continuous information decoding and control, including context-dependent phonological, syntactic and semantic operations (Bulut, 2022; Costafreda et al., 2006; Deniz et al., 2023; Malaia & Newman, 2015; Marklund & Persson, 2012; Mollo et al., 2018; Zhu et al., 2012). The findings also align with studies that identify the IFG as a core region for integrating knowledge during ongoing discourse in reading (Hagoort et al., 2004; Hsu et al., 2015). The results for the AG support previous studies that report its relevance in integrating contextual information during narrative reading (Branzi et al., 2021; Ramanan et al., 2018).

Moreover, the AG is a core region of the DMN and part of the dorsal pathway that includes the IFG. Our findings also suggest that the IFG and AG share a similar neural event timescale, suggesting closer communication and information flow between them. Functional connectivity studies have shown that the prefrontal cortex (e.g. IFG) exhibits stronger connectivity with the DMN (e.g. AG). This connectivity forms a subsystem within the fronto-parietal network, which is involved in introspective processes (Dixon et al., 2018). The short timescale of events suggests that the IFG and AG are involved in integrating semantic information from the written phrase and sentence level, highlighting their role in the rapid construction of information during event segmentation.

Meanwhile, the IFGorb, ATL and PTL were all dominated by relatively longer-lasting events. These three regions are key parts of the ventral pathway for semantic comprehension, are anatomically connected, and share similar functions. Among them, the ATL has been proposed as a hub for semantic representation, facilitating communication between different regions (such as IFGorb and PTL) in the language network according to the Hub-and-spoke theories (Ralph et al., 2017). Furthermore, recent theories emphasize that ATL is the central region for language-derived knowledge representation (Bi, 2021). Using the same dataset, Wehbe et al. also found that the ATL is involved in integrating meaning beyond simple concrete concepts within sentences and narratives (Wehbe et al., 2014). Thus, the ATL is likely a core region for knowledge-based event representation and for facilitating communication with other regions in the reading network. One region that communicates with the ATL is the PTL, which has been recognized for its important role in lexical syntax and semantic processing, as well as in the representation of the thematic and feature-based conceptual knowledge at both the word and sentence levels (Flick et al., 2021; Maran et al., 2022; Matchin et al., 2019; Matchin & Wood, 2020). For example, Matchin et al. (2019) found that the PTL is sensitive to sentence structure versus phrases, as well as to real words compared to non-words. This sensitivity was not observed in the ATL. In addition, Flick et al. (2021) found that both the ATL and PTL are involved in feature-based concept processing, such as the color of an object. However, only the PTL showed sensitivity to the composition of words reflecting an implicit thematic relation (e.g., ‘a robin snake’ is a snake that hunts robins). These findings demonstrate that the PTL might be involved in thematic events based on lexical information. Another region, the IFGorb, is particularly engaged in narrative comprehension (Babajani-Feremi, 2017; Mar, 2011). Previous research has shown that the IFGorb exhibits higher activity when reading an intact story compared to scrambled sentences, reflecting its functional role in maintaining narrative information (Yarkoni et al., 2008). Overall, the findings suggest that the ventral pathway is involved in event segmentation, particularly in processing semantic information that ranges from thematic and knowledge-based concept representation to comprehension.

Interestingly, the MFG exhibited fewer event counts compared to other regions. This difference could be related to the specific role of this area during the reading process. The MFG has been proposed as a site for motor processing, involved in self-generated actions (Henderson et al., 2015; Zhou et al., 2016). For example, the MFG is involved in maintaining the visual serial order of input (Amiez & Petrides, 2007), the linguistic order of events (Munte et al., 2024; Munte et al., 1998; Ye et al., 2011), and reorienting endogenous attentional processes to exogenous stimuli (Japee et al., 2015). It is also involved in pronoun resolution, especially when the pronoun refers to a person compared to a thing, suggesting conceptual rather than syntactic referencing (Hammer et al., 2007). The MFG also responded to the protagonist’s action content (Speer et al., 2009), suggesting involvement in memorizing content. In the current study, words were presented on the screen in an RSVP paradigm during fMRI recording, which minimized participants’ eye movements and increased the working memory load. This may have led to the sustained activation of the MFG for processing serial inputs during reading.

### 4.2 Neural event representation and annotation

Furthermore, we adopted annotations from humans and ChatGPT to interpolate the neural events in the reading network. We investigated whether events in regions within the DMN have a stronger association with annotations under the context of narrative reading. Previous studies revealed that the DMN produces neural events similar to those observed in humans when they process movies or audio stories (Baldassano et al., 2017; Wu et al., 2025). In line with these findings, we found that the AG, the core region of the DMN (Buckner et al., 2008; Raichle, 2015), showed a neural signature of event boundaries that were aligned with human annotations, as well as ChatGPT. This finding confirmed AG’s important role in supporting event segmentation during reading. Together with previous studies on auditory and movie narrative processing, this shows that the AG processes events regardless of the stimulus modality. These findings are consistent with previous studies suggesting that the AG integrates multisensory information into modality-independent concepts (Humphreys et al., 2021; Junker et al., 2024; Kim et al., 2022; Seghier, 2013, 2023), with a particular emphasis on sentence-level topic semantics rather than word-level meaning (Acunzo et al., 2022). Notably, the participants of human annotations were not from the same sample as the fMRI task, and the story was present in the whole text rather than in RSVP format, and still, alignment for most salient boundaries could be detected. This relatively strong association with human annotations might be due to the AG’s significant role in storing and recalling events through direct communication with the hippocampus, a process shared across participants (Chen et al., 2017; Cohen et al., 2022; Liu et al., 2022). Besides, an expanding collection of studies has shown AG’s stronger response to non-linguistic meaning and weaker functional connections with other language-related regions (Fedorenko et al., 2024), so it’s less affected by stimulus presentation type. Overall, our findings indicate that the AG is critical in supporting event segmentation in narrative reading.

The ATL is also a part of the DMN and one of the main semantic processing regions for language (Bemis & Pylkkanen, 2013; Herlin et al., 2021). Our results suggested the ATL reflects the event structure derived from humans, rather than from ChatGPT, which involves less subjective judgment. The literature has shown that the activation of neural populations in the ATL could process domain-general semantic knowledge, such as knowledge of objects, words and facts, especially for social information and personally relevant stimuli (Wong & Gallate, 2012). These results indicate that the neural events in the ATL could encode different semantic information, in contrast to the AG, which is more involved in social and affective features.

We found the inferior frontal areas (e.g. IFG and IFGorb) also showed good alignment with annotations. Recent work demonstrated that these areas play a critical role in the event storage and segmentation control (Baldassano et al., 2017; Geerligs et al., 2022; Lee et al., 2021; Mak et al., 2023; Schapiro et al., 2013; Williams et al., 2022; Yarkoni et al., 2008). For example, Speer et al. (2009) found that the IFG is involved in the event-related time changes and goal redirection of the story’s character. The IFGorb activity also appears to be sensitive to event changes, such as story background information that provides cues for initiating a change in the narrative (Xu et al., 2005). These findings suggest that the involvement of inferior frontal areas in event segmentation might be related to the dimensional features of the situation model.

In summary, our findings suggest that neural events within the DMN and inferior frontal regions within the reading network were associated with annotation. This speaks to a nested interaction of these networks during reading, in line with interactive views of language processing (Fedorenko et al., 2024). The reported HMM approach shows for the first time that these areas play a crucial, yet distinct, role in event segmentation during narrative reading. Our study, along with previous research (Michelmann et al., 2025; Panela et al., 2025), shows that ChatGPT can generate valuable event annotations automatically.

### 4.3 Limitations

An important limitation to consider is the small sample size of the fMRI dataset. The publicly available fMRI dataset includes only eight participants who read one chapter of Harry Potter. Although the dataset spans a lengthy time course (approximately 45 minutes) and has been demonstrated to be reliable and valuable in several studies, it is important to validate our current findings in event segmentation with a larger sample. Additionally, the human annotations were derived from a separate sample, rendering it impossible to study annotation and HMM-based segmentation within the same group, which would be most optimal. A third point is the type of instruction, which was very general for human participants and ChatGPT-3 to avoid a bias towards types of boundaries. As a trade-off, the instruction was open for interpretation of the way to do segmentation. In the future, one could explore segmentation at different situation modality levels (e.g. persons, places, time) and hierarchical organizations within each modality (e.g. fine and coarse events) to further investigate the meaningfulness of HMM-based segmentation results.

## 5. Conclusions

Utilizing the HMM-based event segmentation model, we identified narrative event boundaries within the DMN and reading network, revealing a nested temporal event structure during narrative reading. Neural event boundaries in the IFG and AG were associated with annotations from humans and ChatGPT, suggesting that these regions play a relevant role in situational event segmentation. Taken together, our findings provide evidence of how narrative events are segmented in the reading network.

## Ethics statement

This study was conducted under an approved research line reviewed by the ethics committee of the Faculty of Psychology and Neuroscience, Maastricht University (ERCP205_17_03_2019N-OZL). Informed consent was obtained from all participants who participated in the story annotation task. They received either monetary compensation of 7.5 euros (1 hour) or course credit for their participation.

## CRediT authorship contribution statement

**Huidong Xue:** Conceptualization, Methodology, Software, Investigation, Formal analysis, Data curation, Funding acquisition, Writing - original draft. **Filipp Dokienko:** Methodology, Software, Writing - review & editing. **Francesco Gentile:** Supervision. **Bernadette M. Jansma**: Conceptualization, Methodology, Resources, Supervision, Funding acquisition, Project administration, Writing – original draft, Writing - review & editing.

## Author Approvals

All authors have read and agreed to the published version of the manuscript.

## Acknowledgments

We thank Karolina Niessen and Giovanni Menon for the helpful discussion regarding the Hidden Markov Model, and Elia Formisano for relevant feedback on the first draft of this paper.

## Data and Code Availability Statement

The fMRI dataset is publicly available at https://github.com/anikethjr/brain_syntactic_representations (Reddy & Wehbe, 2021), or https://www.cs.cmu.edu/~fmri/plosone/ (Wehbe et al., 2014). All methods described in this paper are implemented in an open-source Python package called Brain Imaging Analysis Kit (https://brainiak.org/).

